# *Pseudomonas aeruginosa* promotes persistence of *Stenotrophomonas maltophilia* via increased adherence to depolarized respiratory epithelium

**DOI:** 10.1101/2022.06.29.498203

**Authors:** Melissa S. McDaniel, Natalie R. Lindgren, Caitlin E. Billiot, Kristina N. Valladares, Nicholas A. Sumpter, W. Edward Swords

**Author notes:** Communicating author: 1918 University Boulevard, MCLM 818 Birmingham, AL 35294.

## Abstract

*Stenotrophomonas maltophilia* is an emerging opportunistic respiratory pathogen in patients with cystic fibrosis (CF). *S. maltophilia* is frequently observed in polymicrobial infections, and we have previously shown that *Pseudomonas aeruginosa* promotes colonization and persistence of *S. maltophilia* in mouse respiratory infections. In this study, we used host and bacterial RNA sequencing to further define this interaction. To evaluate *S. maltophilia* transcript profiles we used a recently described method for selective capture of bacterial mRNA transcripts with strain specific RNA probes. We found that factors associated with the type IV pilus, including the histidine kinase subunit of a chemotactic two-component signaling system (*chpA*), had increased transcript levels during polymicrobial infection. Using immortalized CF respiratory epithelial cells, we found that infection with *P. aeruginosa* increases adherence of *S. maltophilia*, at least in part due to disruption of epithelial tight junctions. In contrast, an isogenic *S. maltophilia chpA* mutant lacked cooperative adherence to CF epithelia and decreased bacterial burden *in vivo* in polymicrobial infections with *P. aeruginosa.* Similarly, *P. aeruginosa* lacking elastase (*lasB)* did not promote *S. maltophilia* adherence or bacterial colonization and persistence *in vivo*. Based on these results, we conclude that disruption of lung tissue integrity by *P. aeruginosa* promotes adherence of *S. maltophilia* to the lung epithelia in a type IV pilus-dependent manner. These data provide insight into *S. maltophilia* colonization and persistence in patients in later stages of CF disease and may have implications for interactions with other bacterial opportunists.

**IMPORTANCE:** Despite advances in treatment options for patients with cystic fibrosis (CF), complications of bacterial infections remain the greatest driver of morbidity and mortality in this patient population. These infections often involve more than one bacterial pathogen, and our understanding of how inter-species interactions impact disease progression is lacking. Previous work in our lab found that two CF pathogens, *Stenotrophomonas maltophilia* and *Pseudomonas aeruginosa* can cooperatively infect the lung to cause more severe infection. In the present study, we found that infection with *P. aeruginosa* promotes persistence of *S. maltophilia* by interfering with epithelial barrier integrity. Depolarization of the epithelial cell layer by *P. aeruginosa* secreted elastase increased *S. maltophilia* adherence, likely in a type IV pilus-dependent manner. Ultimately, this work sheds light on the molecular mechanisms governing an important polymicrobial interaction seen in pulmonary diseases such as CF.

## INTRODUCTION

*Stenotrophomonas maltophilia* is a Gram-negative bacillus that can be found in a variety of environmental sources, including in hospital tubing and water systems (1–4). As an opportunistic pathogen, *S. maltophilia* is most commonly associated with respiratory infections including ventilator-associated pneumonia (VAP), and chronic airway diseases like cystic fibrosis (CF) (5–8). In the context of CF, detection of *S. maltophilia* in patient sputa has been correlated with worse lung function (9, 10). Whole genome sequencing of *S. maltophilia* has revealed homologs of many known virulence factors including fimbriae, flagella, and type IV pili (11). There is a pressing need for a better definition of factors involved in colonization, persistence and/or virulence of *S. maltophilia*.

*Pseudomonas aeruginosa* is a Gram-negative bacillus that, like *S. maltophilia*, can be found in a variety of environmental contexts. It is an opportunistic pathogen, primarily affecting those with an underlying immunodeficiency or disease, and is a common opportunist observed in patients with CF, where it contributes significantly to morbidity and mortality (8). *P. aeruginosa* has a relatively large genome (∼6.5 Mb), harboring many virulence factors that have been identified and characterized (12). Importantly, *P. aeruginosa* can secrete a number of toxins and extracellular proteases, notably ExoA, elastase, and pyocyanin, that can contribute to lung function decline and can work synergistically to compromise airway barrier integrity (13).

In chronic lung diseases such as CF, infections are often polymicrobial, and inter-species dynamics can play a large role in patient outcomes. Reports indicate that *P. aeruginosa* can cause polymicrobial infections with *S. maltophilia* in patients with CF, VAP, and more recently, hospital acquired pneumonia in patients hospitalized for COVID-19 (14–17). Several *in vitro* studies have suggested mechanisms of cooperativity between *S. maltophilia* and *P. aeruginosa*, including changes in antibiotic tolerance and biofilm formation by *S. maltophilia*, and increased alginate and toxin production by *P. aeruginosa* (18, 19). In previous work, we demonstrated cooperativity between *P. aeruginosa* and *S. maltophilia* during polymicrobial infection in the mouse respiratory tract (20). In this study, intratracheal infection with *S. maltophilia* and *P. aeruginosa* resulted in a significantly higher bacterial burden of *S. maltophilia* in lung homogenate, and a longer time to clearance as compared to mice infected with *S. maltophilia* alone.

In this study, we sought to understand the mechanism by which *P. aeruginosa* promotes colonization with *S. maltophilia.* We used combined bacterial and host RNA-sequencing from murine pulmonary infections with *in vitro* adherence assays on polarized epithelium to elucidate the systems involved in cooperativity between *S. maltophilia* and *P. aeruginosa*. The results indicate that damage to the airway epithelium by *P. aeruginosa* elastase expression promotes increased adherence of *S. maltophilia,* likely via the type IV pilus.

## RESULTS

### Host response to single-species and polymicrobial infection is dominated by *P. aeruginosa*-induced inflammatory response

Our recent work showed a cooperative interaction between two CF pathogens, *S. maltophilia* and *P. aeruginosa,* during murine pulmonary infection wherein the presence of *P. aeruginosa* promotes the persistence of *S. maltophilia.* (Fig. 1A) (20). In order to define the basis for this cooperativity, we first performed host RNA-sequencing analysis (RNA-seq) on whole lung from mice with mono- or polymicrobial infections. Mice were infected intratracheally with *S. maltophilia* K279a (inoculum ∼10^7^ CFU), *P. aeruginosa* mPA08-31 (inoculum ∼10^7^ CFU), or both, before total RNA was collected from the lung, prepared for sequencing, and sequenced at a depth of ∼30 million reads per sample.

**FIG 1.**
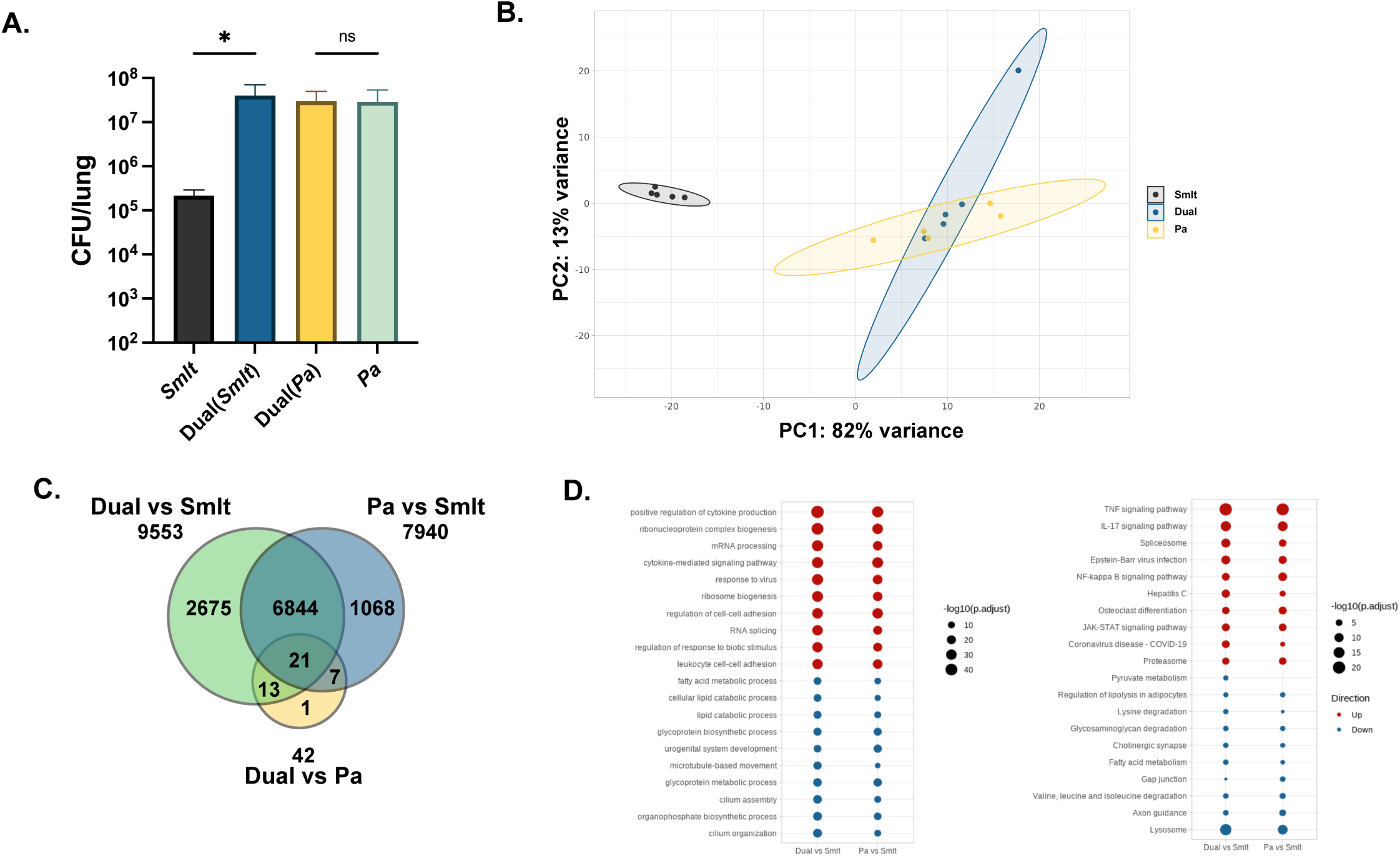
The host inflammatory response following polymicrobial infection is driven by *P. aeruginosa*. BALB/cJ mice were intratracheally infected with ∼10⁷ CFU of *S. maltophilia* K279a and *P. aeruginosa* mPA08-31 alone, and in combination. Groups were euthanized at 24 hours post-infection. A) Bacterial load in lung homogenate was enumerated via viable colony counting on differential medium (mean ± SEM, n = 9-10; Kruskal-Wallis with Dunn’s multiple comparisons, * P<0.05). Using the same infection scheme, whole-lung RNA was extracted and sequenced (n=5). B) Principal component analysis of samples based on RNA-sequencing data, colored by infection group. C) A Venn-diagram depicting the number of significantly differentially expressed genes between groups as determined via differential-expression analysis using DESeq2. D) Pathway analysis of differentially expressed genes performed via clusterProfiler using GeneOntology (biological function) (left) and KEGG pathway databases (right). The top 20 differentially regulated pathways (10 most enriched by positively expressed genes and 10 most enriched by negatively expressed genes) are represented for each comparison.

Principal component analysis of mouse gene expression data showed close clustering of *P. aeruginosa* and polymicrobial infection samples, while *S. maltophilia* infected animals clustered separately from both (Fig. 1B). Differential expression analysis between samples showed that 9,553 transcripts showed differential expression levels between polymicrobial infected mice and mice infected with *S. maltophilia* alone. Similarly, 7,940 transcripts differed between mice infected with *P. aeruginosa* alone and mice infected with *S. maltophilia* alone. Consistent with the principal component analysis, only 42 genes were differentially regulated between mice infected with *P. aeruginosa* alone and polymicrobial infected mice. Of the 9,553 differentially expressed transcripts between polymicrobial infected mice and mice infected with *S. maltophilia* alone, 6,844 were also differentially expressed between mice infected with *P. aeruginosa* alone and mice infected with *S. maltophilia* alone. Only 21 transcripts were differentially expressed in all three comparisons (Fig. 1C).

To determine which biological processes or pathways were affected during infection, we performed pathway enrichment analysis on the list of differentially expressed genes from each comparison. This was performed using ClusterProfiler (21) with both Gene Ontology (GO) biological processes and Kyoto Encyclopedia of Genes and Genomes (KEGG) pathway databases (Fig. 1D). Upregulated genes from the polymicrobial and *P. aeruginosa* infections compared to *S. maltophilia* infection were enriched for a total of 2,206 unique GO terms (1952 and 1918 respectively) and 81 unique KEGG pathways (75 and 68 respectively). The 10 most enriched GO terms among upregulated genes included positive regulation of cytokine production (p_adj_ = 6.71 x 10^-46^, 7.81 x 10^-39^), cytokine mediated signaling pathways (p_adj_ = 1.12 x 10^-40^, 1.76 x 10^-36^), and the regulation of cell-to-cell adhesion (p_adj_ = 1.30 x 10^-31^, 1.68 x 10^-33^).

Of the 10 most enriched KEGG pathways implicated by upregulated genes, we identified known pro-inflammatory pathways including TNF (p_adj_ = 2.99 x 10^-21^, 4.02 x 10^-20^), and IL-17 signaling (p_adj_ = 9.03 x 10^-13^, 8.79 x 10^-13^). The enrichment of these biological processes is consistent with an increase in acute inflammatory response and lung damage during *P. aeruginosa* infection (22). Downregulated genes in these comparisons were enriched for 1,321 unique GO terms (1,108 and 1,073 respectively) and 50 unique KEGG pathways (42 and 32 respectively). Interestingly, both cilium organization and cilium assembly processes were among the 10 most enriched GO terms for downregulated genes (p_adj_ = 1.89 x 10^-20^, 6.45 x 10^-14^), indicating possible disruption of the mucociliary clearance mechanism (23). The 10 most enriched KEGG pathways among downregulated genes highlighted many metabolic processes including amino acid degradation and fatty acid metabolism.

### Selective capture of bacterial mRNA and *in vivo* RNA-sequencing

Traditionally, RNA-sequencing of pathogen transcripts in the lung is difficult due to the overwhelming proportion of host RNA as compared to bacterial RNA. To circumvent this, we employed a previously published method for selective hybridization and capture of bacterial mRNA, previously named pathogen-hybrid capture (PatH-Cap) (24). Strain-specific RNA probe libraries are used to capture pathogen-specific transcripts of interest, allowing for enrichment of bacterial mRNA transcripts and sequencing of the pathogen transcriptome with sufficient coverage, even in the context of RNA extracted from host tissue. We first generated an RNA probe set specific to *S. maltophilia* K279a, comprised of consecutive 100 bp segments covering each annotated open reading frame. The entire sense strand was covered, and a probe was generated for every other 100 bp segment on the antisense strand, as has been shown to increase efficiency of hybridization (24), for a total of 68,704 probes (Fig. S1). Probes were then synthesized as a pool of DNA oligonucleotides by Genscript (Piscataway, NJ), and reverse transcribed to produce a collection of biotinylated RNA “bait” probes.

Total RNA was isolated from lung tissue of mice infected as described above. We then hybridized the bacterial probes to the prepared cDNA libraries, and isolated hybridized transcripts via streptavidin bead-binding. Enriched cDNA pools were then sequenced at a depth of ∼30 million reads per sample (Fig. 2). For the samples from mice infected with *S. maltophilia* in the absence of *P. aeruginosa*, transcript capture resulted in a 697-fold increase in reads mapping to the *S. maltophilia* transcriptome (from 0.01% prior to enrichment, to 6.97% post enrichment). For those samples from mice infected with *S. maltophilia* and *P. aeruginosa*, this increase was 770-fold (from 0.10% to 77.01%) (Table S1).

**FIG 2.**
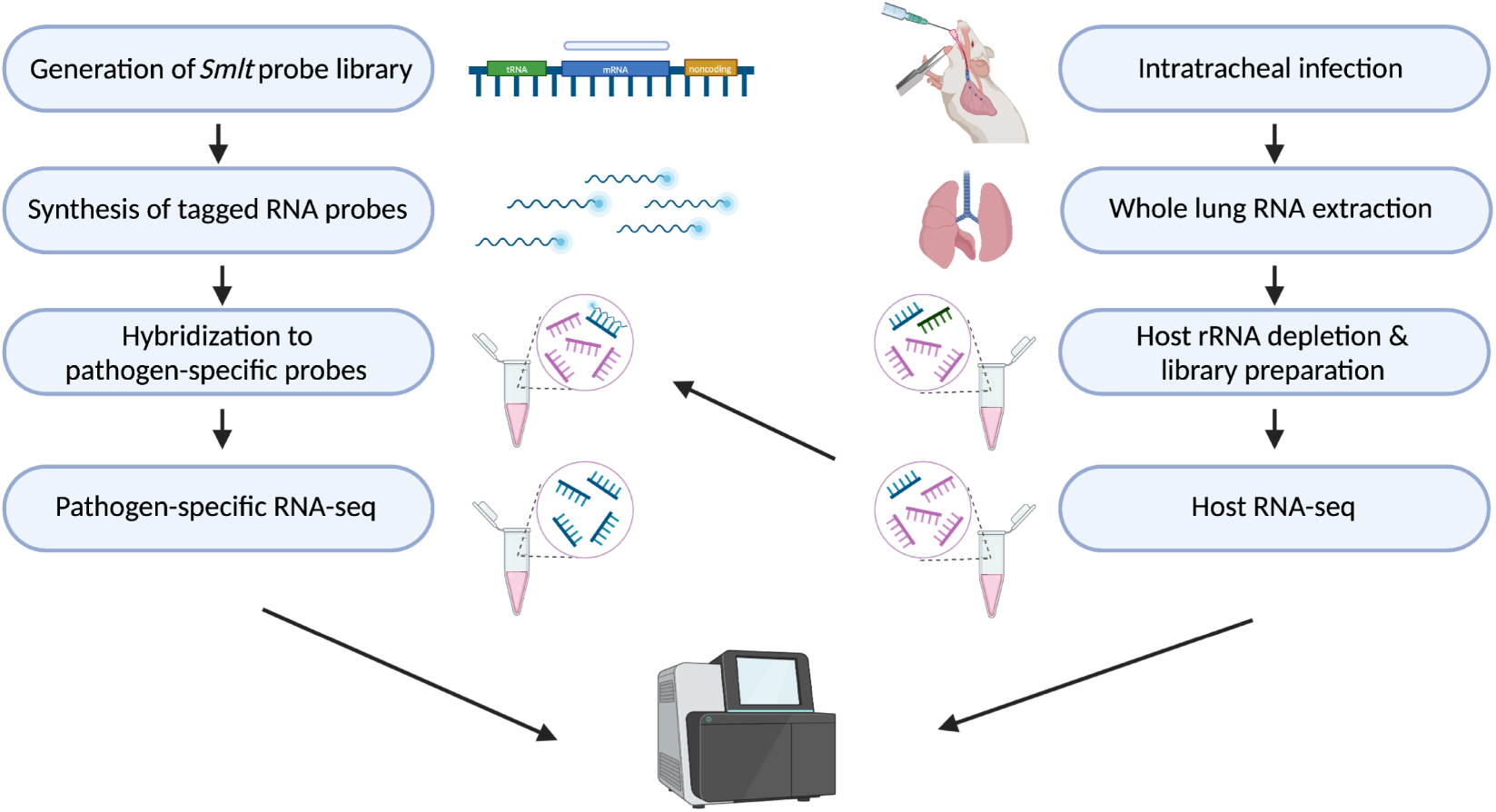
RNA isolation and sequencing methodology. A schematic depicting the workflow of host-pathogen RNA seq using pathogen-hybrid capture (PatH-cap). BALB/cJ mice were intratracheally infected with ∼10⁷ CFU of *S. maltophilia* K279a and *P. aeruginosa* mPA08-31 alone, and in combination, and groups were euthanized at 24 hours post-infection. For host RNA-sequencing, whole-lung RNA was extracted, depleted of host rRNA, and then prepared for sequencing. For bacterial RNA-sequencing, a pathogen-specific probe set was generated to cover coding sequences of *S. maltophilia* K279a. This pool of DNA probes was synthesized, amplified, and then reverse transcribed to create biotinylated RNA probes. Prepared cDNA libraries from the whole lung preparations were hybridized to pathogen-specific probes, and the enriched RNA population was isolated via streptavidin-bead binding. Pathogen-enriched libraries were then sequenced to obtain a bacterial transcript profile.

### Polymicrobial infection increases expression of adherence and chemotaxis-related genes in *S. maltophilia*

Principal component analysis of selective-capture enriched *S. maltophilia* transcript data showed distinct clustering between samples from mice infected with *S. maltophilia* alone and dual species infected samples (Fig. 3A). Differential expression analysis between samples showed 686 *S. maltophilia* genes that are differentially regulated (p_adj_ < 0.05) between these two conditions. To account for disparities in genome coverage between sample groups, we filtered results for genes with detectable transcripts in 2 out of 4 total samples in each group, resulting in a total of 149 differentially expressed genes. Of these, the top 5 significantly upregulated genes included a previously uncharacterized serine protease (*Smlt4395,* p_adj_ = 3.90 x 10^-5^), and two genes involved in type IV pilus biogenesis or regulation, *chpA* (*Smlt3670,* p_adj_ = 4.70 x 10^-5^) and *pilO* (*Smlt3823,* p_adj_ = 2.63 x 10^-4^) (25–27). Interestingly, one of the most significantly downregulated genes during polymicrobial infection was *cheR* (*Smlt2250,* p_adj_ = 2.24 x 10^-4^), a determinant in the regulation of flagellar movement (28), indicating that motility and attachment processes are changing in the context of polymicrobial infection (Fig. 3B). In support of this, of the 19 genes in 3 of the operons predicted to govern type IV pilus biogenesis and regulation in *S. maltophilia*, 12 genes (shown in color) were significantly upregulated during polymicrobial infection in the lung (Fig. 3C). This was not the case for genes involved in fimbriae or flagella regulation and biogenesis. Only 1 of 47 total predicted flagella-related genes were significantly upregulated during polymicrobial infection, and no there was no significant change for *smf-1*, the major protein involved in fimbrial function (Table S2).

**FIG 3.**
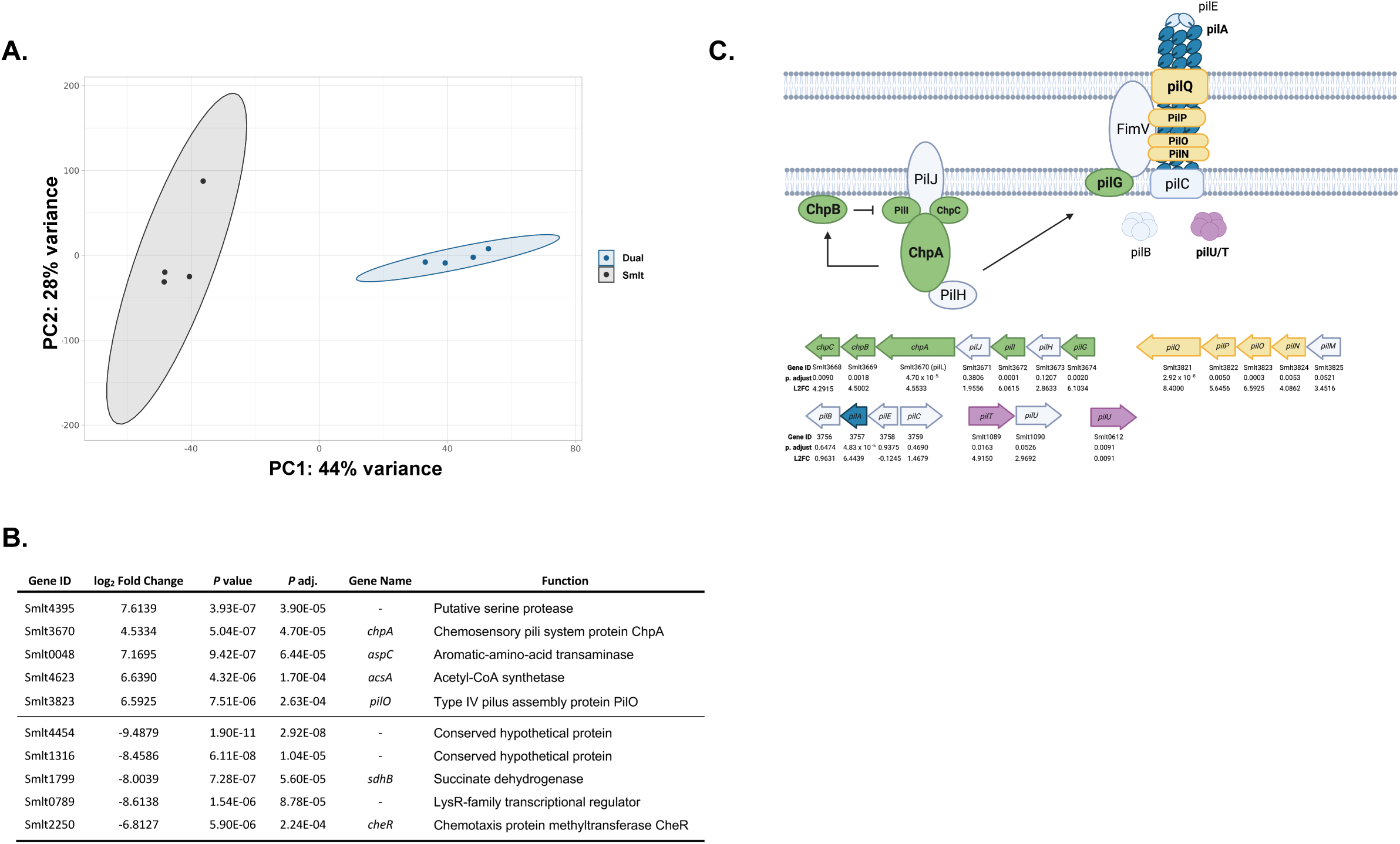
*S. maltophilia* upregulates genes associated with adhesion and chemotaxis in the context of polymicrobial infection. Pathogen-enriched RNA was extracted as detailed above and sequenced (n=4). A) Principal component analysis of samples based on bacterial-specific RNA-sequencing data, colored by infection group. B) Top 5 most significantly up- and down-regulated *S. maltophilia* genes during co-infection in the lung as compared to single-species infection, determined via DESeq2. Genes with reads detected in less than half of the samples for each group were excluded from this analysis. *P* adj. indicates the significance value after multiple testing corrections. C) A schematic depicting the proposed type IV pilus system of *S. maltophilia* (27, 51, 52) and the corresponding differential expression values for each gene. Loci are represented as annotated in *S. maltophilia* K279a (11). Genes significantly upregulated during dual species infection are highlighted in color for each locus.

### Pre-infection with *P. aeruginosa* increases adherence of *S. maltophilia* to polarized epithelia

The increased expression of genes involved in bacterial chemotaxis and adherence, combined with previous reports that exposure of epithelial cells to *P. aeruginosa* can promote *S. maltophilia* adherence (29) prompted us to investigate whether this was a viable mechanism for microbial cooperativity in the lung. To do this, we first polarized immortalized cystic fibrosis bronchial epithelial cells (CFBEs) by culturing them at the air-liquid interface. We then pre-treated the polarized epithelia with *P. aeruginosa* mPA08-31 (MOI = 20) for 2, 4, or 6 hours prior to inoculation with *S. maltophilia* K279a (MOI = 20) for 1 hour. Following this, we evaluated adherence via viable colony counts and confocal microscopy (Fig. 4A). We found that prior infection of cells with *P. aeruginosa* significantly increased the number of adherent *S. maltophilia* (P < 0.0001), with the largest difference occurring at 6 hours post-infection (Fig. 4B) which corresponds with the time point at which the burden of *P. aeruginosa* is the highest (Fig. 4C). Imaging of infected cells via confocal scanning laser microscopy (CSLM) showed more *S. maltophilia* present on cells previously infected with *P. aeruginosa* than on those exposed to cell culture media alone at all time points. We also found that *S. maltophilia* bound to epithelial cells near *P. aeruginosa,* with the largest foci of both bacteria present in cells following preceding infection with *P. aeruginosa* for 6 h. (Fig. 4D).

**FIG 4.**
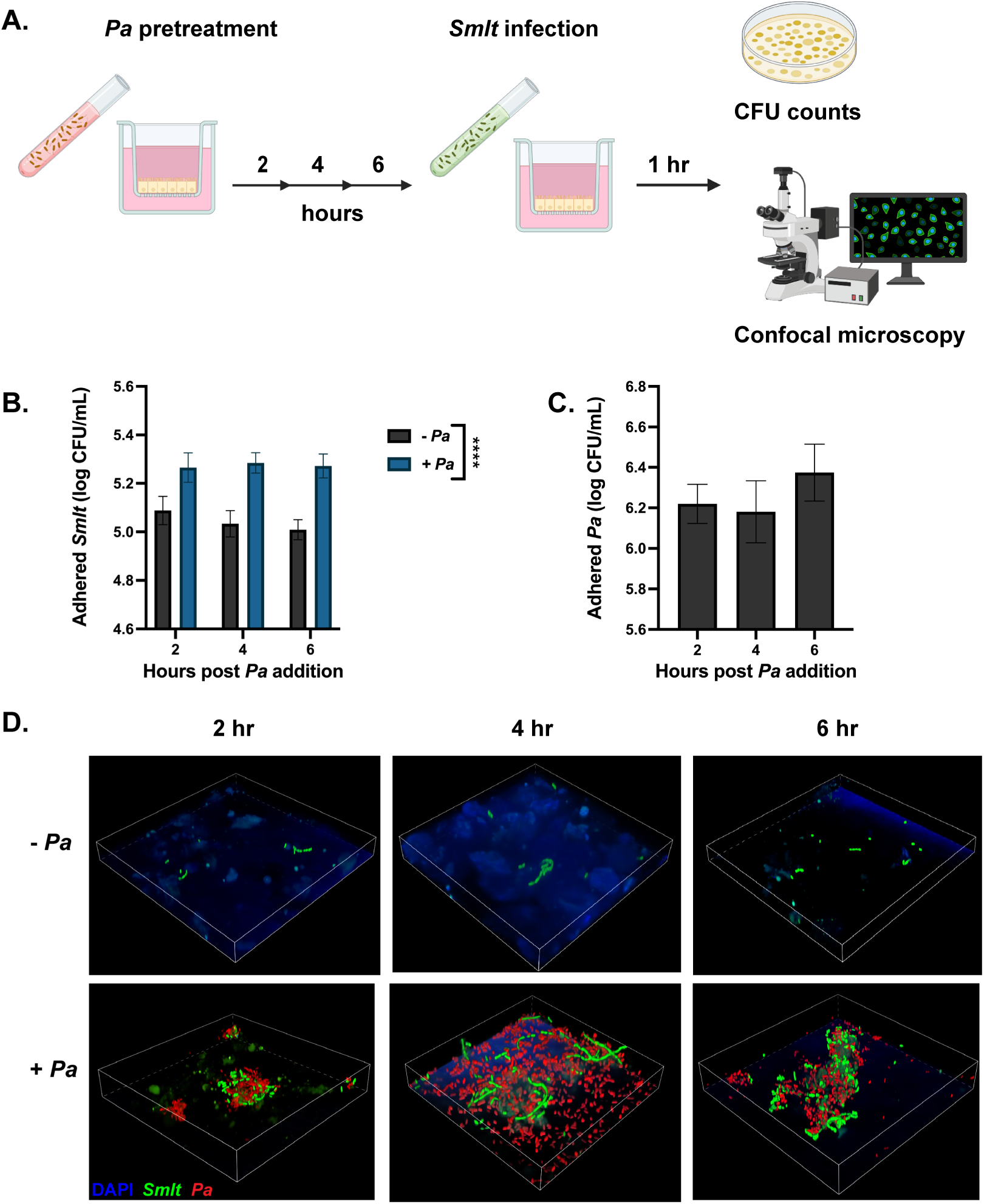
Pre-exposure of epithelial cells to *P. aeruginosa* promotes adherence of *S. maltophilia*. Immortalized cystic fibrosis bronchial epithelial cells (CFBEs) were grown at air-liquid interface until polarized. A) Infection schematic depicting the pre-treatment of with either EMEM or ∼10^6^ CFU of *P. aeruginosa* mPA08-31 for 2, 4, or 6 hours before the addition of ∼10^6^ *S. maltophilia* K279a for 1 hour. B) Viable colony counts of adherent *S. maltophilia* K279a (Mean ± SEM, n = 15 wells. Two-way ANOVA, **** P<0.0001) or C) *P. aeruginosa* mPA08-31 (Mean ± SEM, n = 15 wells). D) Structural composition of polymicrobial foci as evaluated via confocal microscopy. Infections were repeated with *S. maltophilia* K279a (gfp+) and *P. aeruginosa* mPA08-31 (mCherry+), and CFBEs were visualized with DAPI. Polymicrobial foci were imaged at 60X magnification.

### Infection with *P. aeruginosa* promotes adherence of *S. maltophilia* to a polarized epithelium in a *chpA* dependent manner

The most significantly upregulated type IV pilus-related transcript identified in the RNA-seq experiment, *chpA* (*Smlt3670*), was the histidine kinase subunit of a two-component regulatory system characterized in *P. aeruginosa* and known to govern twitching motility (30). To see if this gene impacts cooperativity during polymicrobial infection, we created a clean deletion mutant of *chpA* in *S. maltophilia* K279a (see detailed procedures in Methods). We infected polarized CFBE epithelia with *P. aeruginosa* mPA08-31 for 2 h, 4 h, or 6 h and then added *S. maltophilia* K279a or *S. maltophilia* K279a *chpA* and quantified adhered bacteria via viable colony counts. As shown previously, prior infection with *P. aeruginosa* significantly increases the number of adhered *S. maltophilia* (P < 0.0001). However, *S. maltophilia chpA* had significantly decreased adherence to the CFBE epithelial layer (P < 0.0001), which was unaffected by preceding infection with *P. aeruginosa* (Fig. 5A). These data were confirmed by confocal laser scanning microscopy (CLSM) imaging of infected CFBE epithelial cells. The amount of *S. maltophilia* bound to cells increased when *P. aeruginosa* was present and could be seen adherent to the same regions as large clusters of *P. aeruginosa*. However, *S. maltophilia chpA* had significantly fewer adherent *S. maltophilia*, even in areas with abundant *P. aeruginosa* (Fig. 5B). To test impact of *S. maltophilia chpA* in vivo, we infected mice with both *S. maltophilia* and *S. maltophilia chpA* (inoculum ∼10^7^ CFU) in the presence and absence of *P. aeruginosa* (inoculum ∼10^7^ CFU) for 24 hours. We found that coinfection with *P. aeruginosa* still increased the burden of *S. maltophilia* K279a *chpA* in the lung as compared to single-species infection (Fig. 5C). However, this increase was to a lesser degree than with parental *S. maltophilia*, with a 289-fold increase in *S. maltophilia* burden during coinfection as compared to a 40-fold increase with *S. maltophilia chpA* (Fig. 5D).

**FIG 5.**
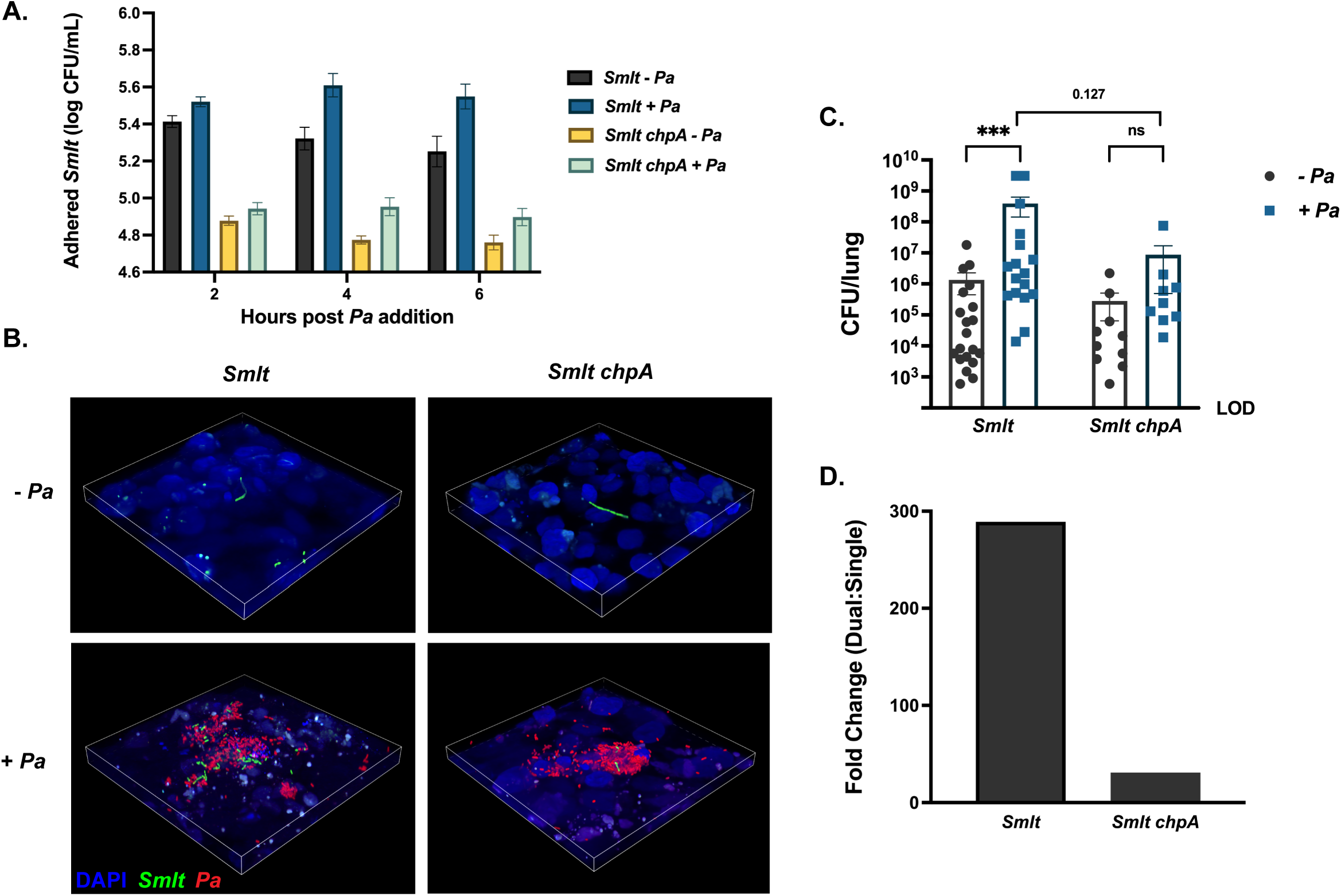
Dysregulation of the type IV pilus abrogates promotion of *S. maltophilia* adherence. Polarized CFBEs were infected ∼10^6^ CFU of *P. aeruginosa* mPA08-31 or vehicle (MEM) for 2 h, 4 h, or 6 h before infection with ∼10^6^ *S. maltophilia* K279a or K279a *chpA* for 1 h. A) Viable colony counts of adherent *S. maltophilia* (Mean ± SEM, n = 9 wells. Two-way ANOVA, **** P<0.0001). B) Structural composition of polymicrobial foci as evaluated via confocal microscopy. Infections were repeated with *S. maltophilia* K279a (gfp+) or K279a *chpA* (gfp+) and *P. aeruginosa* mPA08-31 (mCherry+), and CFBEs were visualized with DAPI. Polymicrobial foci were imaged at 60X magnification. BALB/cJ mice were intratracheally infected with ∼10⁸ CFU of *S. maltophilia* K279a or K279a *chpA* in the presence and absence of *P. aeruginosa* mPA08-31 and were euthanized 24-hours post-infection. C) Bacterial burden in the lung enumerated via viable colony counting from lung homogenate (Mean ± SEM, n = 9-20. Kruskal-Wallis, *** P<0.001). D) Fold change in bacterial counts between single-species and polymicrobial infections.

### Loss of barrier integrity promotes binding of *S. maltophilia* to the bronchial epithelium

*P. aeruginosa* harbors many virulence factors that affect lung barrier integrity during infection, including several secreted proteases (31, 32). Therefore, we hypothesized that breakdown of tight-junctions, and the resulting depolarization of the epithelial cell layer, promotes adherence of *S. maltophilia*. Infection of polarized CFBEs with *P. aeruginosa* (MOI = 20) for 6 hours dramatically decreased organization of occludin-stained tight junctions as compared to cell culture medium alone (Fig. 6A). The transepithelial electrical resistance (TEER), a measurement of monolayer polarity, also decreased significantly over time with the introduction of *P. aeruginosa* (P < 0.0001) (Fig. 6B). To determine if the increase in *S. maltophilia* adherence could be induced by monolayer depolarization in the absence of *P. aeruginosa,* we treated cells with 16 mM EGTA, a calcium chelator that has been shown to delocalize tight-junction proteins including occludin and ZO-1 (33, 34). EGTA treatment for 30 minutes successfully depolarized the epithelial monolayer, with TEER decreasing significantly (P < 0.0001) (Fig. 6C). As expected, significantly more *S. maltophilia* adhered to epithelial cells treated with EGTA as compared to cell culture media controls (P < 0.0001). In contrast, depolarization of the membrane with EGTA did not increase the number of adherent *chpA*-deficient *S. maltophilia* (Fig. 6D). To determine if *S. maltophilia* bound to the specific areas of the cell layer with breakdowns in tight-junction integrity, we stained for the tight junction protein ZO-1 and *S. maltophilia* on cell layers with and without EGTA treatment. Without EGTA, the intercellular tight junctions remained intact and few *S. maltophilia* are present. After pre-treatment with EGTA, the localization of ZO-1 to tight junctions was diminished, consistent with loss of tight junction integrity and depolarization of the epithelia. In these infected cells more adherent *S. maltophilia* were observed, preferentially localized to areas with poor ZO-1 organization (Fig. 6E). Based on these data we conclude that breakdown of tight junctions is sufficient to promote colonization with *S. maltophilia* and that *S. maltophilia* is likely binding to host factors exposed during breakdown of the epithelial barrier rather than to *P. aeruginosa* cellular or biofilm components.

**FIG 6.**
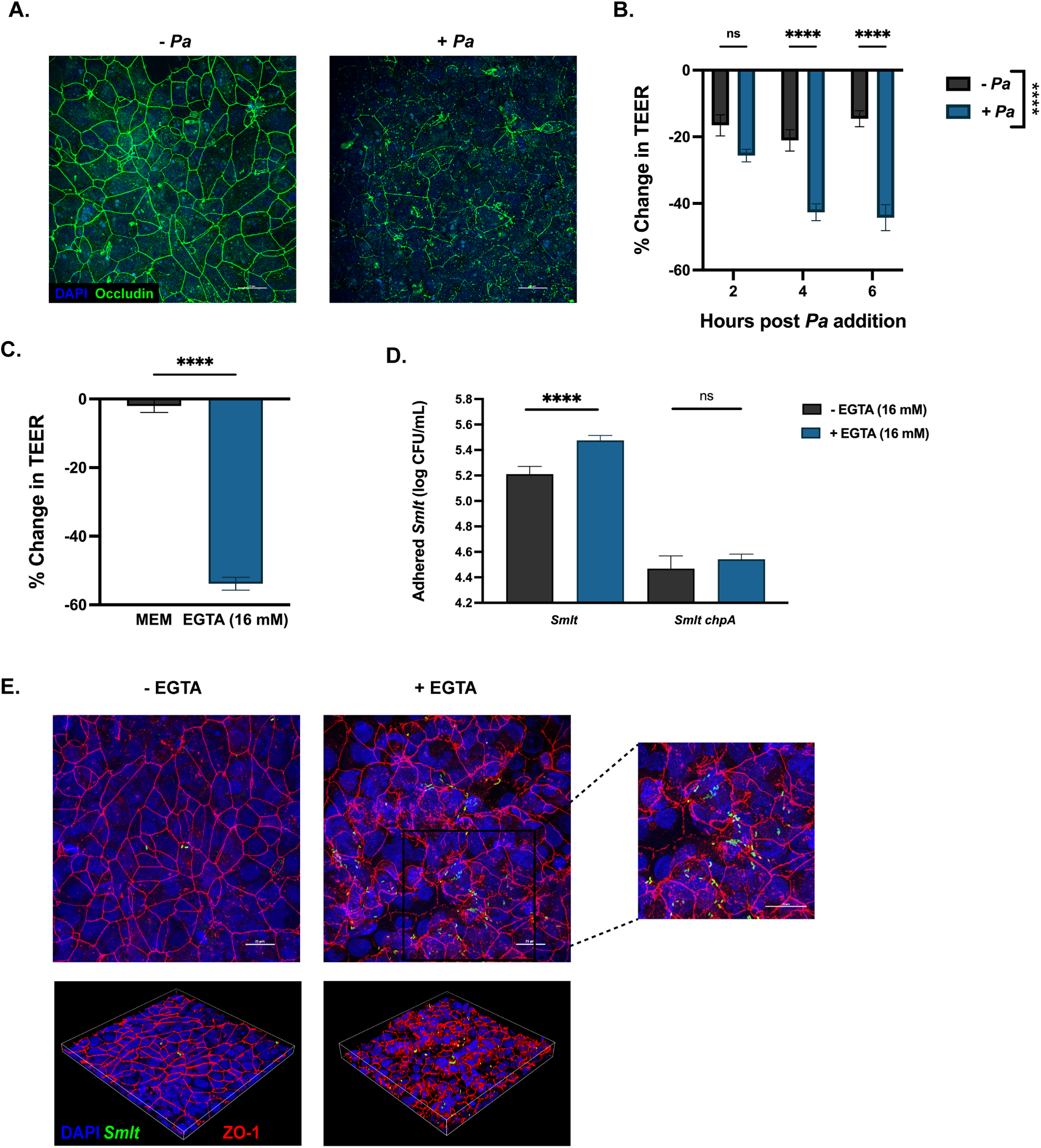
Depolarization of the epithelia is sufficient to promote adherence of *S. maltophilia*. A) Confocal imaging of tight junctions stained via immunfluorescent staining for occludin from cells treated with either EMEM or ∼10^6^ CFU of *P. aeruginosa*. Cells were imaged at 60X magnification. B) Percent change in transepithelial electrical resistance (TEER) across the epithelial membrane after addition of MEM or ∼10^6^ CFU of *P. aeruginosa* (Mean ± SEM, n = 6 wells. Two-way ANOVA, *** P<0.001). C) Percent change in TEER across the epithelial membrane after addition of MEM or 16 mM EGTA for 30 minutes. (Mean ± SEM, n = 20 wells. Unpaired t-test, **** P<0.0001). D) Viable colony counts of adherent *S. maltophilia* K279a and K279a *chpA* on CFBEs after a 30-minute pre-treatment with either MEM or EGTA (16 mM) (Mean ± SEM, n = 12 wells. One-way ANOVA, **** P<0.0001). E) Confocal imaging of CFBEs pre-treated with either MEM (left) or 16 mM EGTA (right) before *S. maltophilia* inoculation. Cells were stained via ZO-1 (red) for tight junctions and *S. maltophilia* via anti-*Smlt* rabbit sera (green) and were imaged at 60X magnification. Inserts were zoomed in 2X for a total magnification of 120X.

### Elastase-mediated damage to the lung epithelium by *P. aeruginosa* increases *S. maltophilia* binding

Production of degradative enzymes by *P. aeruginosa* is an important virulence factor that can interfere with airway barrier integrity and damages host tissue (31, 32, 35). Of these, the secreted protease elastase B is well characterized for its role in pathogenesis. Elastase B is known break down tight junctions, and therefore depolarize epithelial and endothelial cell layers. In combination with toxins secreted by the type III secretion system (T3SS), elastase can also contribute to epithelial invasion and disseminated infection by *P. aeruginosa* (36).

Because we found that tight junction degradation was associated with increased adherence of *S. maltophilia* to host epithelium (Figure 6), we generated an isogenic *P. aeruginosa* elastase deficient mutant (*P. aeruginosa* mPA08-31 *lasB)* which was used to test impact of elastase on epithelial integrity and *S. maltophilia* adherence. We infected polarized CFBE epithelia with *P. aeruginosa* mPA08-31 or *P. aeruginosa* mPA08-31 *lasB,* or mock-infected with cell culture media, for 2 h, 4 h, or 6 h. We then added *S. maltophilia* K279a, quantified adherent bacteria by viable colony counts, and monitored change in TEER over time. After 6 h of incubation *P. aeruginosa* promoted *S. maltophilia* adherence to a greater degree than *P. aeruginosa lasB,* despite no difference in *P. aeruginosa* burden between parental and knockout strains (Fig. 7A, B). Consistent with these results, *P. aeruginosa* decreased TEER across the epithelial monolayer to a greater degree than *P. aeruginosa lasB*, although this was not statistically significant (Fig. 7C). These results were confirmed via confocal imaging of cell monolayers stained for *S. maltophilia* and tight junctions (ZO-1). More *S. maltophilia* was present when cells were pre-infected with *P. aeruginosa* mPA08-31 than when infected with mPA0831 *lasB*. ZO-1 organization was also much better preserved in the cell layer infected with the *lasB* mutant as compared to the parental *P. aeruginosa* strain (Fig. 7D). To evaluate the contribution of elastase *in vivo*, we repeated mouse infection experiments using *P. aeruginosa* and *P. aeruginosa lasB* in conjunction with *S. maltophilia.* The data clearly show that coinfection with *P. aeruginosa* again significantly increased the *S. maltophilia* bacterial colonization and persistence, but there was no such impact in mice coinfected with *S. maltophilia* and *P. aeruginosa lasB* above that of mice infected with *S. maltophilia* alone (Fig. 7E). *P. aeruginosa lasB* resulted in a 3-fold increase in *S. maltophilia* burden during coinfection as compared to a 350-fold increase with the parent strain (Fig. 7F). These results indicate that elastase production by *P. aeruginosa,* and likely the resulting inflammation and lung damage, are necessary factors for the cooperative behavior of these two organisms in the murine lung.

**FIG 7.**
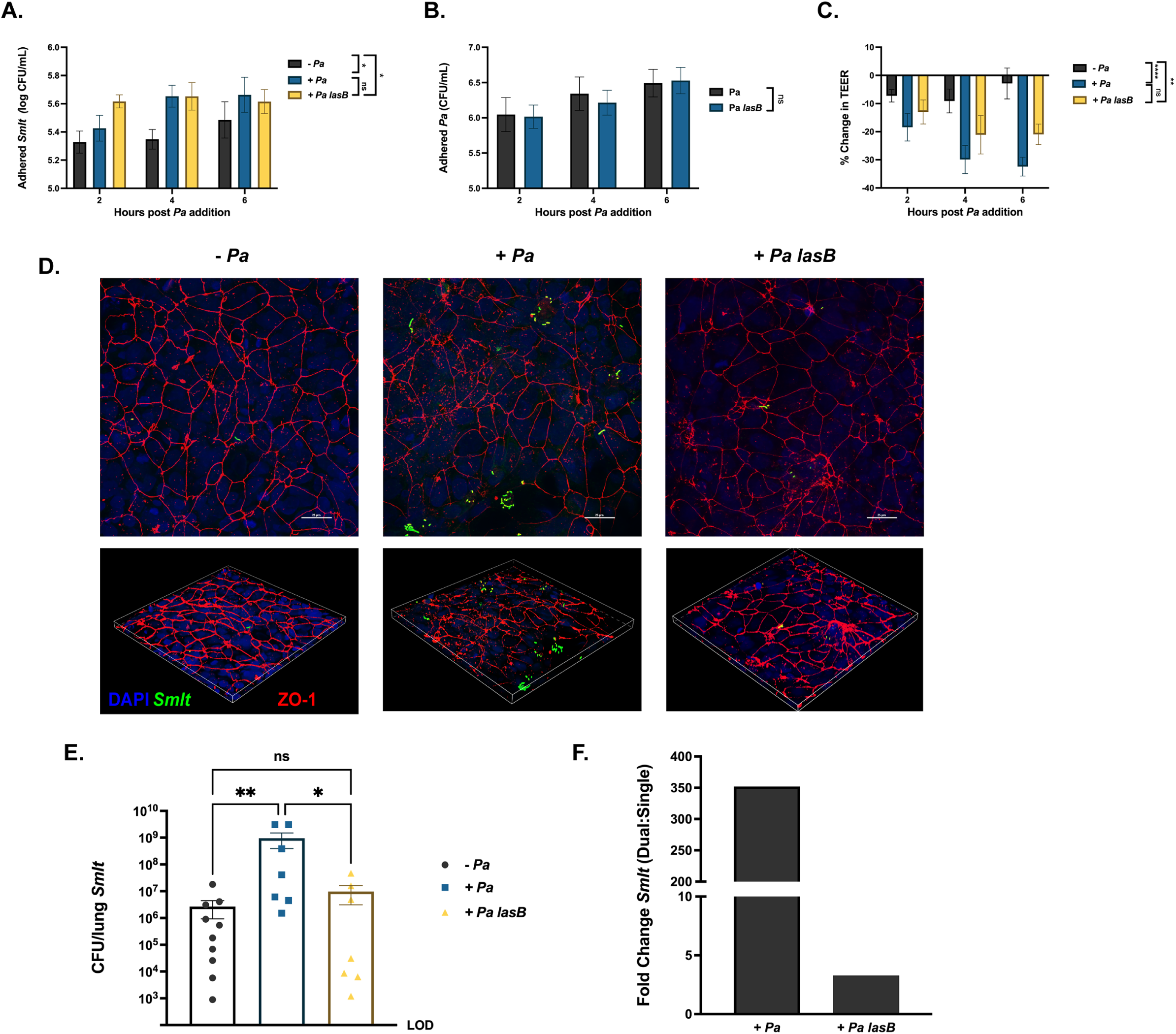
Elastase production by *P. aeruginosa* promotes increased persistence of *S. maltophilia* in the murine lung. Polarized CFBEs were pre-treated with MEM, ∼10^6^ CFU of *P. aeruginosa* mPA08-31 or ∼10^6^ CFU of *P. aeruginosa* mPA08-31 *lasB* for 2, 4, or 6 hours before the addition of *S. maltophilia* K279a for 1 hour. A) Viable colony counts of adherent *S. maltophilia* or B) *P. aeruginosa* (Mean ± SEM, n = 8 wells. Two-way ANOVA, * P<0.05). C) Percent change in TEER across the epithelial membrane after addition of MEM, mPA08-31, or mPA08-31 *lasB* (Mean ± SEM, n = 8 wells. Two-way ANOVA, ** P<0.01, **** P<0.0001). D) Confocal imaging of CFBE41s pre-treated with either WT or elastase deficient *P. aeruginosa* before *S. maltophilia* inoculation. Cells were stained via ZO-1 (red) for tight junctions and *S. maltophilia* via anti-*Smlt* rabbit sera (green) and were imaged at 60X magnification. Inserts were zoomed in 2X for a total magnification of 120X. BALB/cJ mice were intratracheally infected with ∼10⁸ CFU of *S. maltophilia* K279a alone or in the presence *P. aeruginosa* mPA08-31 or mPA08-31 *lasB* and were euthanized 24-hours post-infection. E) Bacterial burden in the lung enumerated via viable colony counting from lung homogenate. (Mean ± SEM, n = 7-10. Kruskal-Wallis, * P<0.05, ** P<0.01). F) Fold change in bacterial counts between single-species and polymicrobial infections.

## DISCUSSION

Respiratory infections have significant impacts on morbidity and mortality in cystic fibrosis and other chronic pulmonary diseases. While many pathogens responsible for these infections have been identified, the advent of culture-independent detection methods has led to an appreciation of the complex ecology of the lung and the impact that inter-species interactions have on patient outcomes. Our prior work showed that colonization and persistence of *S. maltophilia* in the murine lung was significantly increased in polymicrobial infections with *P. aeruginosa*. We also demonstrated that this increased persistence resulted in a higher mortality rate among the infected mice (20). In the present study, we showed that membrane depolarization and lung damage by *P. aeruginosa* mediates increased binding and persistence of *S. maltophilia* in the murine lung.

Previous metrics of virulence indicated that *P. aeruginosa* infection drives the host response during polymicrobial infection, and that the response to *S. maltophilia* is comparatively less severe (20). Our results from RNA-sequencing experiments are concordant with this finding. Principal component analysis of host gene expression suggests that infection with *P. aeruginosa* elicited a similar host response to concurrent infection with both bacterial species. When compared to *S. maltophilia* infection, host genes involved in both cytokine production and cytokine mediated signaling pathways were upregulated in polymicrobial infected mice and mice infected with *P. aeruginosa* alone, indicating a larger magnitude of innate immune response is being mounted when *P. aeruginosa* is present. Although these two organisms share many cellular structures able to elicit a strong immune response (endotoxin, flagella, pili, etc.), the ability of *P. aeruginosa* to damage the lung epithelium through the release of toxins is likely contributing to this response. Genes involved in the cell-to-cell adhesion pathway were also upregulated, indicating a breakdown of barrier integrity.

Host-microbe RNA-seq is a powerful technique that allows for a snapshot of gene expression from both the pathogen and the host in different infection contexts. An important limitation of this technique is the ability to obtain adequate bacterial reads from RNA samples that are overwhelmingly made up of host transcripts. For example, we found that a sequencing depth of ∼30 million reads from lung samples of mice infected with *S. maltophilia* alone yielded on average 0.01% of reads mapping to *S. maltophilia.* To overcome this limitation, we employed a recently published selective mRNA capture technique known as PatH-Cap (pathogen-hybrid capture), which allows for pathogen-specific coding sequences to be enriched from a pool of host and non-coding transcripts (24). For the previously mentioned example that contained 0.01% bacterial reads, application of PatH-Cap increased bacterial reads to 6.97%, a 697-fold increase in relative abundance. PatH-Cap was even more effective with RNA samples from polymicrobial infected mice. The initial bacterial read percentage was 10-fold higher at 0.10%, and the final read percentage was at ∼80%, a 770-fold increase in relative abundance. This illustrates that PatH-Cap is highly dependent on the percentage of input bacterial RNA in the pooled RNA sample. Given sufficient starting material, we have demonstrated that PatH-Cap is a renewable and cost-effective strategy for investigating bacterial RNA expression in disease-relevant contexts, overcoming a major limitation in the bacterial pathogenesis field.

The *in vivo* bacterial RNA-seq results indicated that genes involved in control and biogenesis of the type IV pilus are upregulated in the context of polymicrobial infection in the lung. This included *chpA*, the histidine kinase subunit of a two-component system that, although largely uncharacterized in *S. maltophilia*, is known to regulate twitching motility, mechano/chemotaxis, and cAMP regulation in *P. aeruginosa* (25, 30, 37). Deletion of *chpA* in *S. maltophilia* completely prevented adherence to polarized CFBE cells. Interestingly, previous reports has shown that both flagella and fimbriae of *S. maltophilia* can mediate adherence to epithelial cells (29, 38, 39). However, this work was limited to abiotic surfaces, cell lines not derived from pulmonary epithelium, or non-polarized epithelial layers. Extensive work in *P. aeruginosa* has shown that both flagella and the type IV pilus can mediate adherence to epithelial cells, but show vastly different substrate specificity, with the type IV pilus binding preferentially to host N-glycans on the apical surface of polarized epithelia, and flagella binding preferentially to heparin sulfate proteoglycans on the basal surface (40). It therefore seems likely that *S. maltophilia*, like *P. aeruginosa,* may use several different adherence mechanisms in a context-dependent or redundant manner.

While we know that *S. maltophilia* and *P. aeruginosa* can cause polymicrobial infections, this is certainly not the only risk factor for acquisition of *S. maltophilia.* Decreased lung function, previous antibiotic use, and in-dwelling device all predispose to *S. maltophilia* infection (41). The results herein, although important for understanding a polymicrobial interaction, also clarify how a damaged lung environment might be sufficient to promote *S. maltophilia* binding, as would be the case in patients with late-stage CF disease. EGTA experiments demonstrate that depolarization of the cell monolayer in the absence of *P. aeruginosa* is sufficient to induce increased *S. maltophilia* adherence. Damage and subsequent depolarization of epithelial membranes can expose receptors or ligands that allow for more effective pathogen binding and the role of previous lung damage in establishment of future infection is well characterized for many other pathogens (42, 43). There is certainly the possibility for other variables impacting *S. maltophilia* colonization or persistence, including changes in cellular immune response or nutrient availability. An important next step in this work would be the identification of the host factor responsible for *S. maltophilia* adherence, particularly in the context of a damaged lung environment (44).

With the advent of effective modulator therapies in CF, patient variables including the rapid decline in lung function and persistent inflammation associated with chronic infection with chronic infections may be changing. As with most opportunistic airway infections, CF related respiratory infections are polymicrobial in nature and there is potential for interspecies influences on colonization and persistence in the respiratory tract. With regard to *S. maltophilia*, our findings demonstrate that preceding or coinciding infections may be major determinants of infection outcomes.

## MATERIALS AND METHODS

### Strains and growth conditions

*S. maltophilia* K279a is a widely used model strain with a fully annotated genome sequence, originally isolated from a patient with bacteremia in the UK (11); this strain and the *S. maltophilia* K279a-GFP derivative were provided by M. Herman (Kansas State University). *P. aeruginosa* mPA08-31 was originally isolated from the sputum of a patient with CF and was provided by S. Birket (University of Alabama at Birmingham). *P. aeruginosa* mPA08-31-mCherry+ was constructed by transforming parent strains with plasmid pUCP19+mCherry provided by D. Wozniak (Ohio State University). All strains were routinely cultured on Luria Bertani (LB) agar (Difco) or in LB broth. *S. maltophilia* strains were streaked for colony isolation before inoculating into LB broth and shaking overnight at 30°C, 200 rpm. *P. aeruginosa* strains were streaked for colony isolation before inoculating into LB broth and shaking overnight at 37°C, 200 rpm.

### Mouse respiratory infections

BALB/cJ mice (8-10 weeks old) were obtained from Jackson laboratories (Bar Harbor, ME). Mice were anesthetized with isoflurane and intratracheally infected with either *S. maltophilia, P. aeruginosa,* or both (∼10^7^ CFU each in 100 µL PBS). Mice were euthanized 24 hours post-infection and the left lung of each mouse was harvested and homogenized in 500 µL of sterile PBS for viable plate counting. Homogenate from single-species infections was serially diluted in PBS and plated on LB to obtain viable CFU counts. Homogenate from polymicrobial infections were plated on M9 minimal medium (45) to enumerate *P. aeruginosa* and LB agar containing gentamicin (50 µg/mL) to enumerate *S. maltophilia.* All samples from polymicrobial infections were also plated on LB for total bacterial counts. All mouse infection protocols were approved by the UAB Institutional Animal Care and Use Committees.

### RNA library preparation

For RNA isolation from the lung, BALB/cJ mice (8-10 weeks old) were obtained from Jackson laboratories (Bar Harbor, ME). Mice were anesthetized with isoflurane and intratracheally infected with either *S. maltophilia, P. aeruginosa,* or both (∼10^7^ CFU each in 100 µL PBS). Mice were euthanized 24 hours post-infection, and lungs were inflated with RNA-later to preserve RNA integrity. Whole lungs were homogenized, and cells were lysed by bead beating (0.1 mm silica) in Trizol reagent (Invitrogen), and a full lung RNA extraction was performing using a standard protocol (46). Extracted RNA samples were sent to GENEWIZ (South Plainfield, NJ) for DNase treatment, host and bacterial rRNA depletion (Ribo-Zero Gold rRNA Removal Kit (Epidemiology), Illumina), and library preparation using standard protocols. For bacterial RNA-seq, single-end directional samples were DNase treated (DNase I, NEB) using a standard protocol, before being run through a second Trizol extraction to purify the sample of enzyme. Clean RNA was first rRNA depleted using the NEBNext rRNA depletion kit (Human/Mouse/Rat) (NEB) before being prepared for sequencing using the NEBNext Ultra II Directional RNA Library Prep Kit (NEB) and tagged for multiplexing via the NE Next Multiplex Oligos for Illumina (NEB).

### Pathogen-Hybrid Capture

A pathogen-specific probe list for *S. maltophilia* was generated using previously published methods (24). Briefly, 100 bp probes were generated to cover annotated coding sequences of *S. maltophilia*, with 15 bp spacer sequences added on either end to allow for amplification of the probe library.

5’ SPACER : ATCGCACCAGCGTGT

3’ SPACER: CACTGCGGCTCCTCA

Probes completely covered the sense strand of each gene and were also generated to tile every other 100 bp of the antisense strand for a total of 68,704 probe sequences Figure S1). Probes with significant homology to the mouse genome or to bacterial non-coding RNAs (P <0.05 with BLAST analysis) were manually removed from the list. The resulting oligo pool was synthesized by Genscript (Piscataway, NJ). Before hybridization, DNA oligos were amplified and then reverse transcribed (MEGAshortscript T7 Transcription Kit, Ambion), with added biotin-16-UTP (Roche) to generate biotinylated RNA probes.

Prepared cDNA libraries were hybridized to the generated pathogen-specific RNA probes using previously described methods (24) with a few modifications. Briefly, the synthesized probes and the prepared cDNA library were incubated in hybridization buffer for 24 hours at 68°C. Blocking primers were modified for compatibility with NEB multiplexing primers and a 3’ ddc’ modification was added to maintain barcoding.

Hybridization primer FWD:

AATGATACGGCGACCACCGAGATCTACACTCTTTCCCTACACGACGCTCTTCCGATCT/3ddC/

Hybridization primer REV:

CAAGCAGAAGACGGCATACGAGATNNNNNNNNGTGACTGGAGTTCAGACGTGTGC TCTTCCGATCT/3ddC/

Mouse cot-1 DNA was also swapped for human cot-1 DNA in the hybridization buffer to account for host differences. Hybridized cDNA was isolated via streptavidin beads and then eluted. A diagnostic qPCR (Kapa library quantification kit, Roche) was used to determine the appropriate number of amplification cycles, and enriched samples were amplified using universal primers that maintained sample barcoding before sequencing. All primers used in these experiments are detailed in the supplemental material (Table S3).

### Sequencing, Alignment and Analysis

For host RNA-seq, paired-end strand-specific RNA sequencing was performed using an Illumina HiSeq2 with ∼25,000,000 reads per sample. Reads were trimmed with Trim Galore! (v. 0.4.4) and Cutadapt (v. 1.9.1) and evaluated for quality with FastQC. Reads were aligned to the mouse transcriptome generated from Ensembl gene annotations (build GRCm39/mm39) using the STAR aligner (v. 2.7.3a). Read counts were obtained via Subread FeatureCounts and differential expression analysis was performed with DESeq2, with a ^*p*-value cutoff of < 0.01. Pathway analysis was performed using clusterProfiler (v. 4.4.1) with Gene Ontology (biological processes) and KEGG pathway databases.

Single-end strand-specific sequencing was performed on samples enriched for pathogen-specific RNA via hybridization (∼30,000,000 reads/sample) via Illumina NextSeq500. These reads were again trimmed with Trim Galore! (v. 0.4.4) and Cutadapt (v. 1.9.1) and evaluated for quality with FastQC. Reads were aligned to the published genome of *S. maltophilia* K279a using the STAR aligner (v. 2.7.3a). Read counts were obtained via Subread FeatureCounts and differential expression analysis was performed with DESeq2. Final analysis of significantly up- and down-regulated genes from *S. maltophilia* in the context of polymicrobial infection was restricted to those genes with transcripts detected in at least 2/4 samples from each group.

### Cell culture

Cystic fibrosis bronchial epithelial cells (CFBE41o-) cells, henceforth referred to as CFBEs, are an immortalized human bronchial epithelial cell line homologous for the F508del mutation in CFTR (47) and were propagated from low-passage liquid nitrogen stocks in the Cell Model and Evaluation core laboratory in the UAB Center for Cystic Fibrosis Research. Cells were routinely cultured in minimal essential medium (MEM, Corning) with 10% fetal bovine serum, and were polarized by seeding at a density of ∼10^6^ cells/well on the apical surface of transwells (0.4 μm, Corning) and growing at 37°C for 7 days, before removing the apical media and growing for an additional 7 days at air-liquid interface. Polarization of the epithelial membranes was confirmed via transepithelial electrical resistance measurements performed via EVOM^2^ Volt/Ohm Meter (World Precision Instruments).

### Adherence assays

To measure the adherence of *S. maltophilia* to CFBEs after prior infection with *P. aeruginosa,* cells were inoculated with ∼ 10^6^ CFUs (MOI = 20) of *P. aeruginosa* mPA08-31 in MEM (no FBS). The media on the basal side of the chamber was also replaced with FBS-free medium before incubation. Bacteria were incubated on the cells for 2 h, 4 h, or 6 h before being removed from the apical chamber. Cells were then inoculated with ∼ 10^6^ CFUs (MOI = 20) of *S. maltophilia* K279a and incubated for an hour. Cells were washed twice with sterile PBS before being scraped from the transwell membrane, diluted, and plated on differential medium to enumerate the bacterial burden. TEER was measured for each well both before infection and at the end of *P. aeruginosa* mPA-0831 infection at each time point specified.

For EGTA exposure experiments, cells were treated apically with plain MEM or MEM with 16 mM EGTA 30 minutes at 37°C. Media was then removed and cells were inoculated with ∼ 10^6^ CFUs (MOI = 20) of *S. maltophilia* and incubated for an hour. Bacteria were enumerated by plate count as described above.

### Immunofluorescence Staining and Confocal Microscopy

For imaging of polymicrobial infections, cell layers were infected as described above with *P. aeruginosa* mPA08-31 (mCherry+) and *S. maltophilia* K279a (gfp+) or K279a *chpA* (gfp+). Cells were fixed with 4% paraformaldehyde overnight at 4°C. Cells were then rehydrated with PBS and stained with DAPI. Filters were mounted with ProLong Diamond Antifade (Invitrogen) and were imaged using z-stacks via confocal laser scanning microscopy (CLSM) using a 60X objective.

For imaging of tight-junctions, cells were stained via immunofluorescence. Cells were fixed as previously described, with a few modifications (48). In brief, cells were fixed in 1:1 acetone methanol at 20°C for 10 minutes before rehydrating in TBS. Cell layers were blocked with TBS + 3% BSA for 30 minutes before staining. Cells were incubated with primary antibody (rabbit polyclonal α-occludin, or donkey α-goat ZO-1 for tight junctions, and polyclonal α-*Smlt* rabbit sera cross adsorbed against *P. aeruginosa* for staining of *S. maltophilia*) for 1 hour at room temperature. Filters were then washed and incubated with secondary antibody for 1 hour at room temperature, before being stained with DAPI and mounted with ProLong Diamond Antifade (Invitrogen). Filters were again imaged using z-stacks via confocal laser scanning microscopy (CLSM) using a 60X objective. For insets, images were zoomed in 2X for a total magnification of 120X.

CLSM was performed using a Nikon-A1R HD25 Confocal Laser Microscope (Nikon, Tokyo, Japan). Images were acquired and processed using the NIS-elements 5.0 software.

### Bacterial deletion mutants

An unmarked isogenic deletion mutant of Smlt3670 (*chpA*) in *S. maltophilia* was produced via two-step homologous recombination as has been previously described for *P. aeruginosa* (49). DNA fragments of 500-1000bp upstream and downstream of each gene were inserted in pEX18Tc using the Gibson Assembly Cloning Kit (NEB) using standard protocols from the manufacturer. Plasmids were transformed into *E. coli* DH5α before introduction into *S. maltophilia* K279a via triparental conjugation with helper strain PRK2013 as previously described (50). Clean deletion was confirmed by PCR amplification of the designated region. The unmarked deletion mutant of *lasB* in *P. aeruginosa* mPA08-31 was produced using the methods detailed above, but with a pEX18Gm backbone. For each mutant in *S. maltophilia* or *P. aeruginosa,* at least two independently derived mutants were evaluated and whole genome sequencing was performed showing a lack of secondary mutations.

### Statistical analyses

Unless otherwise noted, graphs represent sample means ± SEM. For non-parametric analyses, differences between groups were analyzed by Kruskal-Wallis test with the uncorrected Dunn’s test for multiple comparisons. For normally distributed data sets (as determined by Shapiro-Wilk normality test) a one-way ANOVA was used with Tukey’s multiple comparisons test. For analyses with more than one factor, a Two-way ANOVA was used. All statistical tests were performed using Graphpad Prism 9 (San Diego, CA).

## DATA AVAILABILITY

Sequencing data generated for all samples included in this study are deposited in the NCBI Sequence Read Archive under the BioProject ID PRJNA853083. Accession numbers for individual sample sequencing read libraries are provided in the supplementary information.

## ACKNOWLEDGMENTS

This work was supported by grants from the Cystic Fibrosis Foundation (CFFSWORDS1810 and CFFSWORDS20G0) awarded to W.E.S. as well as NIH P30 center grant DK072482 awarded to Dr. Steve Rowe. M.S.M. was supported by an NHBLI T32 UAB pre-doctoral training program in lung diseases (T32HL134640, W.E.S. PI) and as a trainee on the UAB CF Research and Development Program (CFF-ROWE19RO).

The authors thank Shawn Williams and Robert Grabski at the UAB High Resolution Imaging Facility for their assistance with the Nikon A1 Confocal microscope and imaging analysis. We also thank Cristina Penaranda (Harvard University), Dr. Michael Crowley (University of Alabama at Birmingham) and the UAB Heflin Genetics core for helping with sequencing, experimental design, and for lending their expertise with regards to the PatH-Cap and RNA sequencing experiments. Megan Kiedrowski provided valuable feedback on the manuscript during revision. We Dr. Bill Benjamin (University of Alabama at Birmingham), Dr. Daniel Wozniak (Ohio State University), Dr. Jessica Scoffield (The University of Alabama at Birmingham), and Dr. Susan Birket (The University of Alabama at Birmingham) provided bacterial strains for this study.

